# Targeted CRISPR-Cas9 screening identifies transcription factor network controlling murine haemato-endothelial fate commitment

**DOI:** 10.1101/2024.01.14.575582

**Authors:** Michael Teske, Tobias Wertheimer, Stefan Butz, Pascale Zwicky, Izaskun Mallona, Ulrich Elling, Christophe Lancrin, Burkhard Becher, Ana Rita Grosso, Tuncay Baubec, Nina Schmolka

## Abstract

Haematopoiesis is a tightly coordinated process that forms and maintains all blood cells. During development blood generation begins in the yolk sac with the differentiation of haemato-endothelial mesoderm giving rise to haematopoietic progenitors. Which molecular regulators are crucial for haemato-endothelial mesoderm formation remains unclear and has not been studied in an unbiased way. Here we employ a mouse embryonic stem cell model that recapitulates embryonic blood development and perform targeted CRISPR-Cas9 knock out screens focusing on transcription factors and chromatin regulators. Focusing on the transition of primitive towards haematoendothelial mesoderm we identified the known master regulator Etv2 and novel transcription factors including Smad1, Ldb1, Six4 and Zbtb7b acting as crucial drivers or repressors of mesodermal commitment. Our transcriptome analysis highlights that each factor has a precise impact on the gene expression signature of the developing mesoderm resulting in the formation of mesodermal subsets with a defined lineage differentiation bias. Our study reveals novel molecular pathways governing mesodermal development crucial to allow endothelial and haematopoietic lineage specification.

## Introduction

Cellular identities result from a combination of genetic and chromatin-based information that ultimately determines individual gene expression programmes. Lineage commitment is achieved by a stepwise acquisition of cell fate changes starting from pluripotent cells of the embryo. While genetic information encodes transcription factors (TFs) that drive progenitor fate and differentiation, chromatin-based information shapes the accessibility of TFs to DNA, providing a heritable chromatin landscape that can direct or reinforce lineage-specific transcriptional programmes. The regulation of early blood specification in the embryo is not fully explored. Blood development is a precisely controlled process that in the mouse initiates at around embryonic day 7 in the so-called blood island of the yolk sac. Blood cells develop from a mesodermal precursor with both haematopoietic and endothelial developmental potential, the haemato-endothelial mesoderm or haemangioblast. Haemato-endothelial mesoderm gives rise to the first wave of blood formation generating primitive erythrocytes and initiates at the same time definitive haematopoiesis by differentiation to haemogenic endothelia. Subsequently, haemogenic endothelia where the endothelial-to-haematopoietic transition (EHT) occurs, gives rise to the second wave of blood formation where mainly erythro-myeloid progenitors (EMPs) are formed. Only a few molecular regulators that impact mesodermal commitment towards a haemato-endothelial fate have been identified, including the best-known master transcription factor ETV2. ETV2 is crucial for the transition of the multipotent primitive mesoderm towards haemato-endothelial mesoderm. While loss of function of ETV2 results in a complete differentiation block of both lineages and early lethality^1,2^ ectopic expression of ETV2 enables fibroblasts to change to the endothelial or haematopoietic fate^2–4^. A recent study employing hypomorph ETV2 mutants highlights a differential dose-dependency for lineage commitment, with the haematopoietic lineage commitment being more sensitive to reduced ETV2 dosage than the endothelial lineage^5^. Several molecular regulators including Brachyury and EOMES are known to be crucial for the initial stages of pluripotent cells to commit to the mesodermal lineage and lie upstream of the gene regulatory network dependent of ETV2^6^. Additionally, we know which factors are needed for blood commitment after haemato-endothelial differentiation has occured including TAL1, RUNX1 and GATA1. Still, until today we lack a largescale interrogation which regulators cooperate to allow the transition of primitive mesoderm with a broad developmental potential to haemato-endothelial mesoderm giving rise to both haematopoietic and endothelial lineages. A comprehensive identification of factors that specify haematopoietic fate during mesodermal development not only provides valuable insights into lineage specification and reprogramming but is essential for the generation of haematopoietic progenitors from pluripotent stem cells for regenerative medicine. In our study we therefore focus on the transition of primitive mesoderm towards haemato-endothelial mesoderm by utilizing FLK1 and PDGFRα surface expression marking this transition. We employed a mouse embryonic stem cell differentiation model that recapitulates embryonic haematopoiesis in the yolk sac^7–9^ and combined this with large-scale CRISPR-Cas9 KO screens focusing on transcription factors and chromatin regulators. We confirmed Etv2 as a master regulator driving primitive towards haematoendothelial mesoderm commitment and in addition, identified novel TFs namely Ldb1, Smad1 and Six4 that are equally needed to allow this transition. We further identify Zbtb7b as a repressor of haemato-endothelial lineage specification, as cells with loss of Zbtb7b function commit more efficiently towards haemato-endothelial mesoderm compared to wild type cells. Additionally, our transcriptomic analysis of the newly defined core TFs shows that they impact distinctly on the gene expression program in primitive mesoderm. Upon loss of function of these core TFs, primitive mesoderm is formed with a defined lineage differentiation bias and distinct properties to generate haematopoietic and endothelial cells. Collectively, by employing large-scale CRISPR screening we identify novel molecular regulators impacting on haematopoietic-endothelial lineage commitment.

## Results

### In depth characterisation of the developmental route to haemato-endothelial mesoderm

To conduct large-scale perturbation experiments and unravel the roles of individual TF and chromatin-regulators facilitating the transition from primitive mesoderm to haemato-endothelial mesoderm, we used a mouse ESC differentiation model that faithfully recapitulates embryonic haematopoiesis (Supplementary Fig 1a). Towards this, we first performed an in-depth phenotypic and molecular characterisation of the ESC based differentiation model. ESC commit towards mesoderm during embryoid body (EB) formation and FLK1 and PDGFRα are known to distinguish distinct differentiation stages: uncommitted cells are negative for both markers (DN), paraxial mesoderm is single positive for FLK1-PDGFRα+ (P_SP), primitive or uncommitted mesoderm is positive for both FLK1+ PDGFRα+ (DP) and haemato-endothelial mesoderm or haemangioblasts are single positive for FLK1+ PDGFRα- (F_SP)^10,11^. In our time course analysis of FLK1 and PDGFRα expression mesodermal commitment started at day 3 when DN cells and P_SP cells were detected and reached a peak at day 5 with the highest frequency of DP/ primitive mesoderm that progresses to F_SP cells / haemato-endothelial mesoderm (Fig. 1a,b). To characterise the distinct mesodermal population and to confirm their developmental trajectory, we performed gene expression analysis employing both single cell and bulk RNA-seq at day 5 when all four populations are presents (DN, P_SP, DP, F_SP). Using our scRNAseq data we performed an RNA velocity analysis and constructed the developmental trajectory from DNèP_SPèDPèF_SP mesodermal populations with the appropriate gene expression patterns (Fig1c, Supplementary Fig. 1b, c). This is in line with previous reports employing ESC differentiation models and with the kinetics of FLK1 and PDGFRα expression in our EB cultures. To further validate the use of FLK1 and PDGFRα as differentiation markers from ESC to haemato-endothelial mesoderm, we preformed bulk-RNAseq on DN, P_SP, DP and F_SP populations at day 5. Using principal component analysis the four populations could be clearly separated (Supp. Fig 1d) and we confirmed the expression of relevant marker genes for distinct developmental stages: pluripotency genes are expressed in the DN population, early mesodermal markers in P_SP and DP, and haemato-endothelial genes in the F_SP (Supp. Fig 1e).

**Figure 1:**
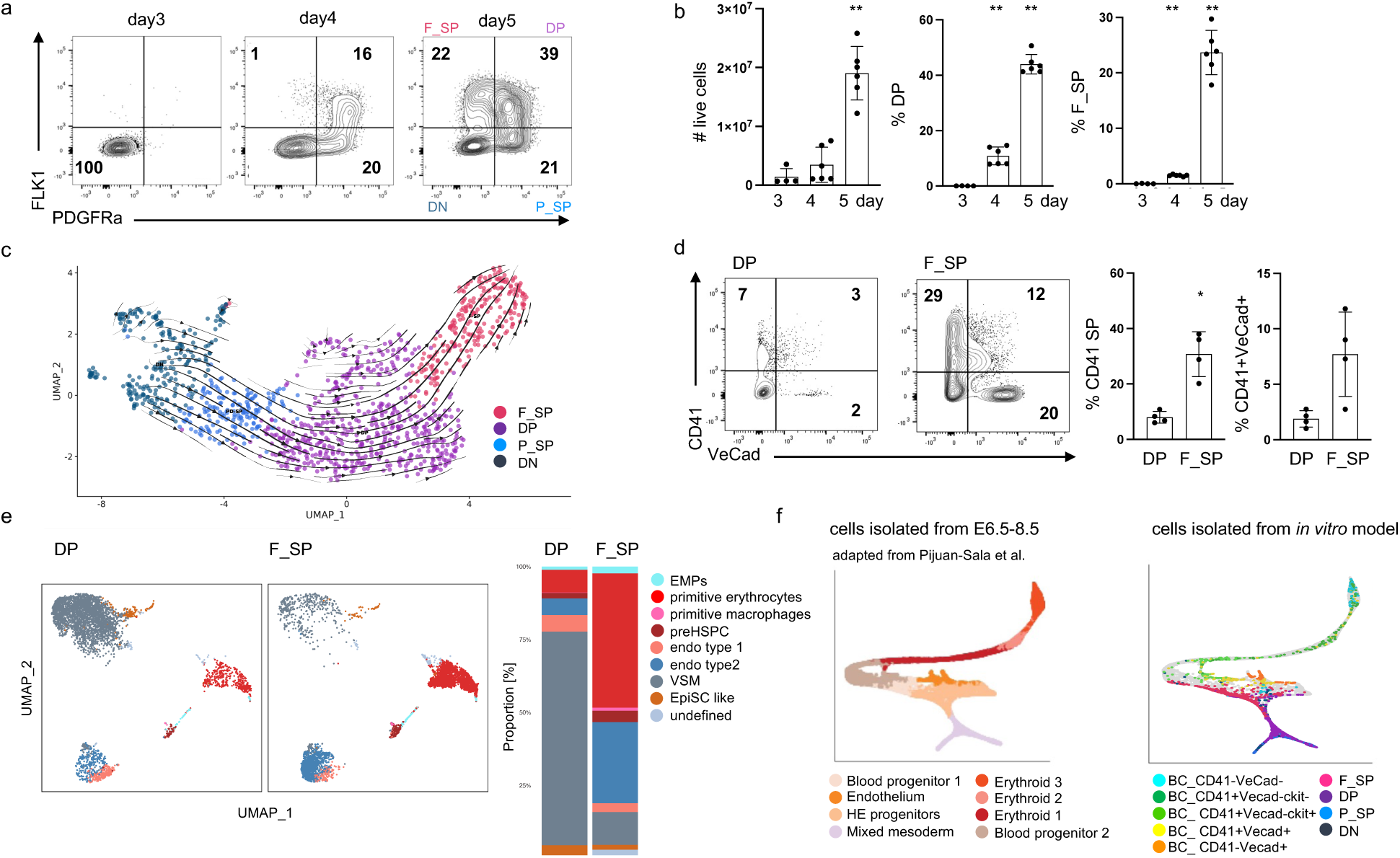
In-depth analysis of the murine ESC-derived differentiation model towards the haemato-endothelial cell fate. **a** Representative flow cytometry analysis of time course analysis of FLK1 and PDGFRα surface expression in WT cells. **b** (left) Number of live cells and (middle) percentage of FLK1+/ PDGFRα + (DP) cells and (right) FLK1+/ PDGFRα - (F_SP) cells in EB cultures over time. **c** UMAP of 1221 cells FACSorted from EB cultures at day5 coloured by cell surface marker expression. **d** Representative flow cytometry analysis of CD41 and VeCadherin (VeCad) surface expression and (left) percentage of CD41+/VeCad- and (right) CD41+/VeCad+ cells in blast culture started from DP or F_SP cells. **e** (right) UMAP with overlaid FlowSOM clustering of 50’000 cells and (left) barchart depicting frequency of cellular subsets identified by FlowSOM clustering in blast culture started from DP or F_SP cells. **f** (left) Force-directed graph layout of cells isolated from E6.5–E8.5 embryos associated with the blood/endothelial lineage (adapted from^15^). Plot highlights various cell types that are generated along the haemato-endothelial lineage trajectory as mesoderm differentiates into endothelial and haematopoietic cells. (right) Mapping of *in vitro* generated cells isolated from EB and blast cultures. Each data point (in b and d) represents an individual experiment (b) n= 6, (d) n=4. Error bars represent mean +/− SD. *P < 0.05 and **P < 0.005, p-values were calculated using an unpaired, two-tailed t test (Mann-Whitney).

To test the developmental potential of DP and F_SP cells we isolated them by FACS sorting and re-plated them in an assay named haemangioblast culture. During this culture blast colony forming cells (BL-CFCs) generate colonies containing both hematopoietic and endothelial progeny and also vascular smooth muscle (VSM)^12,13^. CD41 (first haematopoietic marker) and VE-cadherin (VeCad, endothelial marker) expression is used to distinguish haematopoietic and endothelial fate conversion: haematopoietic (CD41^+^), endothelial (VeCad^+^), preHSPC (positive for both CD41 and VeCad) and VSM cells (negative for CD41 and VeCad)^14^. As expected, DP cells have a reduced capacity to form haematopoietic and endothelial lineages compared to the F_SP population that generates a higher frequency of haematopoietic and endothelial lineages at the expense of vascular smooth muscle cells (Fig. 1d). To get a higher resolution on the lineage potential of the two different mesodermal populations, we established a high-dimensional flow cytometry analysis using 23 surface marker and performed unsupervised clustering to define 9 major clusters regarding their surface expression patterns. Compared to DP cells, F_SP cells formed at higher frequency haematopoietic and endothelial lineages including primitive and definitive blood subsets and endothelial type 1 and type 2 clusters, whereas DP cells have a higher potential to form VSM (Fig.1e, Supp. Fig1f). Furthermore, to underline the relevance of our ESC differentiation model we mapped our *in vitro* generated cells of both stages (EB culture and blast culture) onto a mouse gastrulation atlas of cells isolated from E6.5–E8.5 embryos^15^. As a result, 39.1% of the *in vitro* generated cells from our differentiation model could be projected onto the haemato-endothelial trajectory graph (Fig. 1f). Taken together our single-cell and bulk RNA-seq profiles confirms that FLK1 and PDGFRα are the appropriate markers to distinguish mesodermal populations and that our EB differentiation model faithfully recapitulates early mesodermal commitment and subsequent blood differentiation via an endothelial-to-haematopoietic transition.

### CRISPR screens identify core transcription factors of haemato-endothelial mesoderm commitment

Having defined the developmental route giving rise to F_SP (haemato-endothelial mesoderm) we performed loss-of-function CRISPR screening to identify factors regulating the generation of this population. We first established the CRISPR screen workflow using a library against transcription factors and chromatin regulators targeting 4077 genes by sets of 5 individual guide RNAs (EpiTF library) (Fig. 2a). We transduced the sgRNA library containing a GFP reporter into an ESC line constitutively expressing Cas9 (ref^16^) to ensure optimal KO efficiency. 2 days after transduction, sgRNA expressing ESCs were sorted (based on GFP expression) and sgRNA^+^ ESCs were used to initiate EB differentiation. At day 5 we collected the four distinct mesodermal populations (DN, P_SP, DP, F_SP) using FLK1 and PDGFRα expression (Fig. 2a). We recovered and sequenced gRNAs from the mesodermal populations and a sample collected at the beginning of the EB cultures (T0) for three independent screens and performed sgRNA detection and ranking of individual genes using MAGeCK. To discover which genes regulate the transition from primitive mesoderm towards haemato-endothelial mesoderm, we compared sgRNA presentation in DP versus F_SP populations and generated a ranked candidate gene list by generating the rank mean of the three individual screens (Fig. 2b, c). We identified hits where their targeting sgRNA abundance was either depleted or enriched in F_SP population versus DP population with high statistical ranking, indicating that those genes act as potential drivers or repressors for this lineage transition, respectively (Fig.2b, c). By comparing the top 200 depleted genes we identified 10 common overlaps in the three independent CRISPR screens (Fig. 2d): *Smad1*, *Etv2*, *Ep300*, *Rps27a*, *Mis18a*, *Ldb1*, *Rpa1*, *Med18*, *Six4*, *Hira*. For the top 200 enriched genes we identified 4 overlaps (Fig. 2e): *Cop1*, *Zbtb7b*, *Zic3*, *Npm1*. Interestingly, all overlapping factors displayed a high screen performance and were among the top20 depleted/enriched genes, respectively. As we additionally performed the sgRNA detection in the populations T0, DN and P_SP we compared if our potential hits have differences in the guide abundance (cpm reads) specifically in the transition from DP towards F_SP (Fig. 2f, Supp Fig.2a). This was the case for the 14 overlaps that we detected in the individual screen. Importantly, we detected ETV2, the best-known TF regulating haemato-endothelial mesoderm commitment on rank2, which serves as a positive control for our screening set-up.

**Figure 2:**
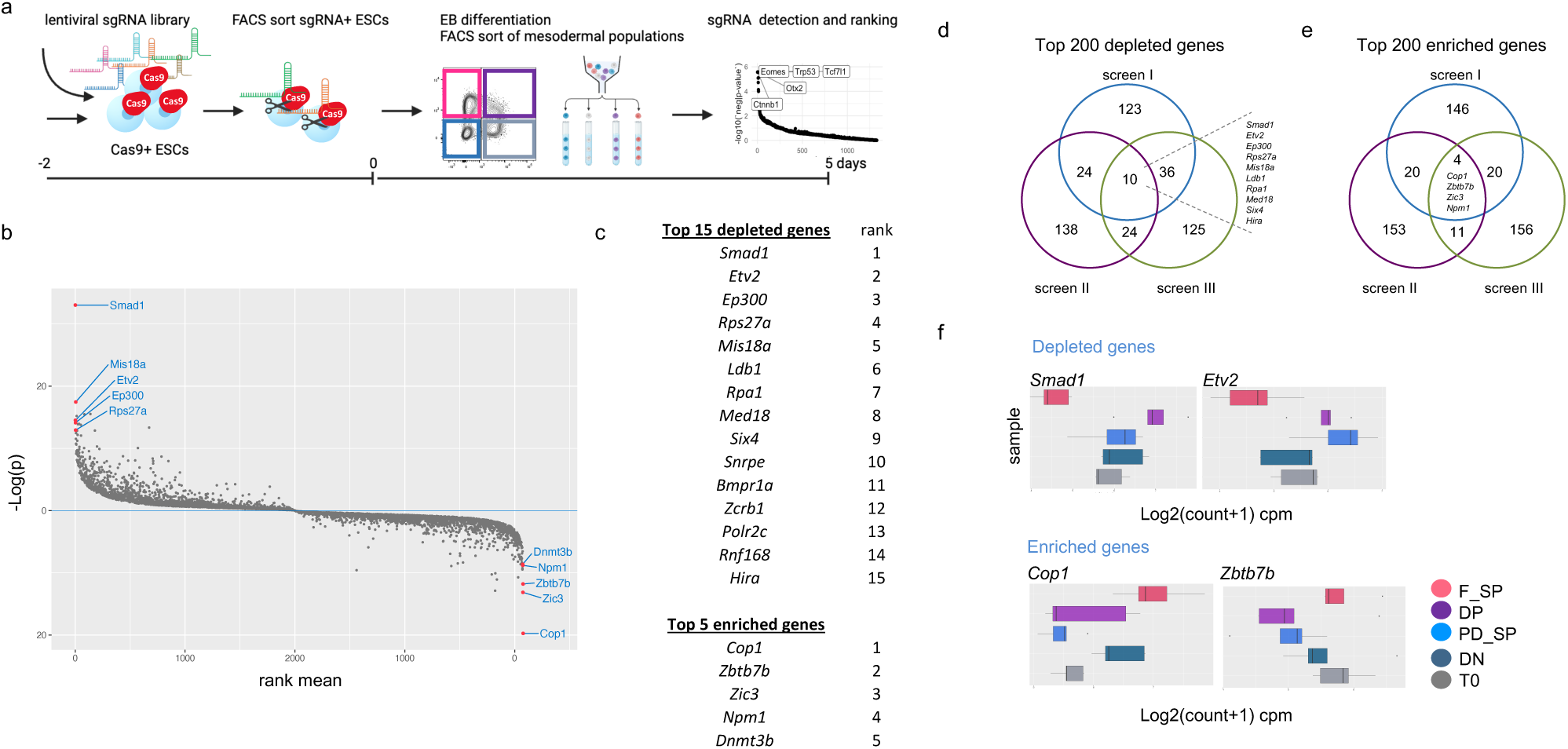
Pooled targeted CRISPR-Cas9 screens identify drivers and repressors of haemato-endothelial mesoderm differentiation. **a** Schematics of pooled targeted CRISPR-Cas9 screens in EB cultures. **b** sgRNA ranking and **c** candidate gene list for the transition of primitive mesoderm (FLK1+ PDGFRα+). towards haemato-endothelial mesoderm (FLK1+ PDGFRα-). **d** Overlaps of top 200 depleted genes in three individual CRISPR screens. **e** Overlaps of top 200 depleted genes in three individual CRISPR screens. (**f**) Representative count per million (CPM) values of sgRNA abundance at T0 and mesodermal populations of top 2 depleted and enriched genes: *Smad1*, *Etv2*, *Cop1* and *Zbtb7b*.

### Validating Hit Genes that promote or repress haemato-endothelial mesoderm commitment

To validate whether the hit genes regulate haemato-endothelial lineage commitment and for manageability of experimental validation, we applied 3 criteria to select genes for validation: Top 20 factors for both the depleted and enriched genes that are additionally common in the three independent screens (among top 200 hits) and have a p-value <0.001. This resulted in a potential candidate list of 13 genes (10 of the depleted genes and 3 of enriched genes) (Supp. Fig. 3a). For each candidate, we cloned independent KO ESC lines using a targeting strategy using two sgRNAs and performed EB differentiation experiments to score their ability to form F_SP cells. Importantly, we generated at least three independent KO lines for each candidate to ensure that their phenotype is driven by the gene KO and compared them to Ctrls transfected with non-targeting guides. For three genes, Mis18a (kinetochore protein), Rps27a (ribosomal protein) and Rpa1 (cell cycle protein) we were not able to generate viable KO ESC lines pointing to an essential role of those genes and we had to exclude those genes from further analysis. Of the 7 hits of the depleted genes 4 candidates resulted in a significant decrease in the frequency of F_SP cells: *Smad1* (rank1), *Etv2* (rank2), *Ldb1* (rank6) and *Six4* (rank 9) (Fig.3a,b). To understand if our candidate genes had a general impact on cellular differentiation, we also analysed the frequency of total FLK1+ (including both DP and F_SP populations) and calculated the ratio of F_SP/ total FLK cells. Importantly, our 4 validated candidates generated similar amounts of total FLK1+ cells compared to the control (Supp. Fig.3b) and had a significant decrease in the ratio of F_SP/ total FLK cells (Fig.3c) pointing to a specific role of those gene for the transition of DP towards F_SP and not an overall defect on mesodermal commitment. Furthermore, no significant differences of total live cells were detected in our validated candidates (Fig. 3d). To our surprise, the gene KOs for Ldb1, Smad1 and Six4 had an equally strong effect on arresting DP cells to further differentiate towards F_SP, compared to the known TF Etv2. Of the 3 hits of the enriched genes we validated 1 hit: Zbtb7b (rank2). EB cultures of Zbtb7b KO cells generated F_SP cells at a significantly higher frequency but resulted in comparable amount of total FLK1+ cells and live counts to the Ctrls (Fig.3e-h, Supp.Fig 3c). These results suggest that Zbtb7b acts specifically as a repressor of haemato-endothelial commitment. Making use of our scRNAseq dataset we analysed the gene expression of the validated candidates along the mesodermal differentiation trajectory. In agreement with previous reports *Etv2* shows a peak of expression at the transition from DP towards F_SP and it not expressed at earlier stages. *Smad 1* gene expression peaks earlier than Etv2 at the beginning of DP development but is broadly expressed in all populations. *Ldb1*, *Six4* and *Zbtb7b* are equally expressed in the different mesodermal populations, with Ldb1 showing the highest and Zbtb7b the lowest level of expression. This data underlines that our candidate genes, except for *Etv2*, would have been difficult to identify by employing a differential gene expression analysis between DP and F_SP cells.

**Figure 3:**
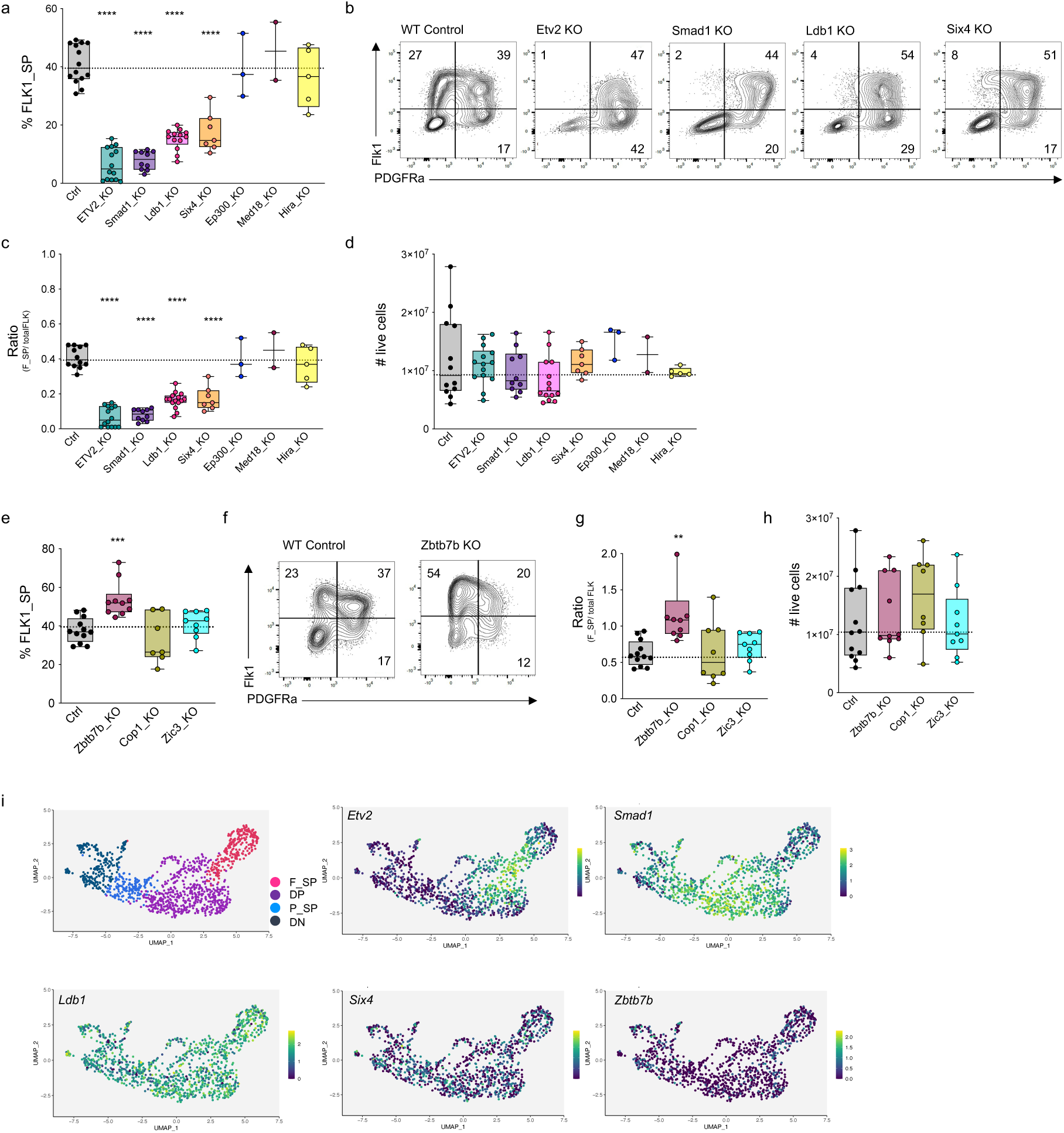
Validation of candidate genes regulating haemato-endothelial mesoderm differentiation. **a, e** percentage of F_SP cells in EB cultures of indicated KOs. **b, f** Representative flow cytometry analysis of FLK1 and PDGFRα surface expression in EB cultures of indicated KOs. **c**,**g** Ratio of F_SP/ total FLK in EB cultures of indicated KOs. **d, h** Number of live cells in EB cultures of indicated KOs. Each data point (in a, c-d, e, g–h) represents an individually generated KO. For each cell line 3 independent KO clones were analysed, n = 4 biological replicates. Error bars represent mean +/− SD. **P < 0.005, ***P < 0.0005, and ****P < 0.0001, p-values were calculated using a one-way ANOVA multiple comparisons analysis. **i** Expression of validated candidate genes in mesodermal populations analysed by scRNA-seq.

Collectively, by applying strict criteria to select genes for validation we were able to validate our hits with a high efficiency. We confirmed the critical role of the already known master regulator Etv2 to enable F_SP cell differentiation and identified novel candidates, Smad1, Ldb1, Six4 and Zbtb7b that act as drivers and repressors of F_SP cell commitment.

### Distinct gene expression signatures in primitive mesoderm of target gene KOs

After establishing the regulatory roles of Etv2, Smad1, Ldb1, Six4 and Zbtb7b in the transition from DP to F_SP, we next wanted to obtain a better insight into the gene expression perturbations in the respective KO lines. Therefore, we performed RNA-seq at the EB stage from FACS sorted DP which are the precursor stage of F_SP cells and equally present in all KO lines and compared them to Ctrl samples transfected with non-targeting guides. The comparison of the global gene expression profiles by principal component analysis indicates similarities between the Ctrl DP cells and DP cells from Etv2 KO, Ldb1 KO and Six4 KO (Fig.4a). The cluster of these cell lines is separated from DP of Smad1 KOs (Fig. 4a). In agreement with this, our differential gene expression analysis between Ctrl and Smad1 KO cells revealed drastic changes in gene expression with 683 genes upregulated and 1313 genes downregulated in the Smad1 KO DP cells (Supp. Fig 4a), whereas the other KOs of Etv2 (114 up / 66 down), Ldb1 (189 up / 330 down) and Six4 (5 up / 92 down) showed lower numbers of differential expressed genes (DEGs) (Supp. Fig. 4a). Surprisingly, minimal overlaps in DEGs were observed when comparing the different KO conditions (Fig. 4b). Gene set enrichment analysis (GSEA) revealed distinct biological processes affected by the different KOs. For Etv2 KO, as expected, we detected several negatively enriched terms related to early blood development including vasculature development, embryonic haematopoiesis and erythroid as well as myeloid development, which was in agreement with the downregulation of known genes critical for blood development such as Tal1, Sox7, Gata1, Gata2, Myc (Fig. 4c, Supp. Fig 4a). The analysis of Smad1 KO revealed a downregulation of terms associated with vasculature development and smooth muscle differentiation, alongside a wealth of terms related to early embryonic development. Interestingly, in the Smad1 KO terms related to neuron fate specification were enriched and not in any other analysed KO (Fig. 4c, d) . Ldb1 KO GO terms were broad, with downregulated terms focusing on stem cell differentiation as well as embryonic organ and head development (Supp. Fig. 4b). Interestingly, no hemopoiesis or erythrocyte differentiation-related terms were enriched below the adjusted p-value cut-off (Fig. 4c). For Six4 KO, we detected several negatively enriched terms related to early blood development similar to Etv2 KO cells (Fig.4c). The vasculature development is significantly downregulated, driven by Six4-DEGs such as Cdh5, Kit, Gli3, Smarca2 or Tgfbr3. In addition, the embryonic hemopoiesis term is depleted with core enrichment genes including Kit, Gata2 and Vegfa, of which only Kit is a Six4 KO DEG (Supp. Fig 4a). In three KOs, Etv2, Smad 1 and Six 4, we detected a significant down-regulation of endothelial cell fate (Fig. 4d).

**Figure 4:**
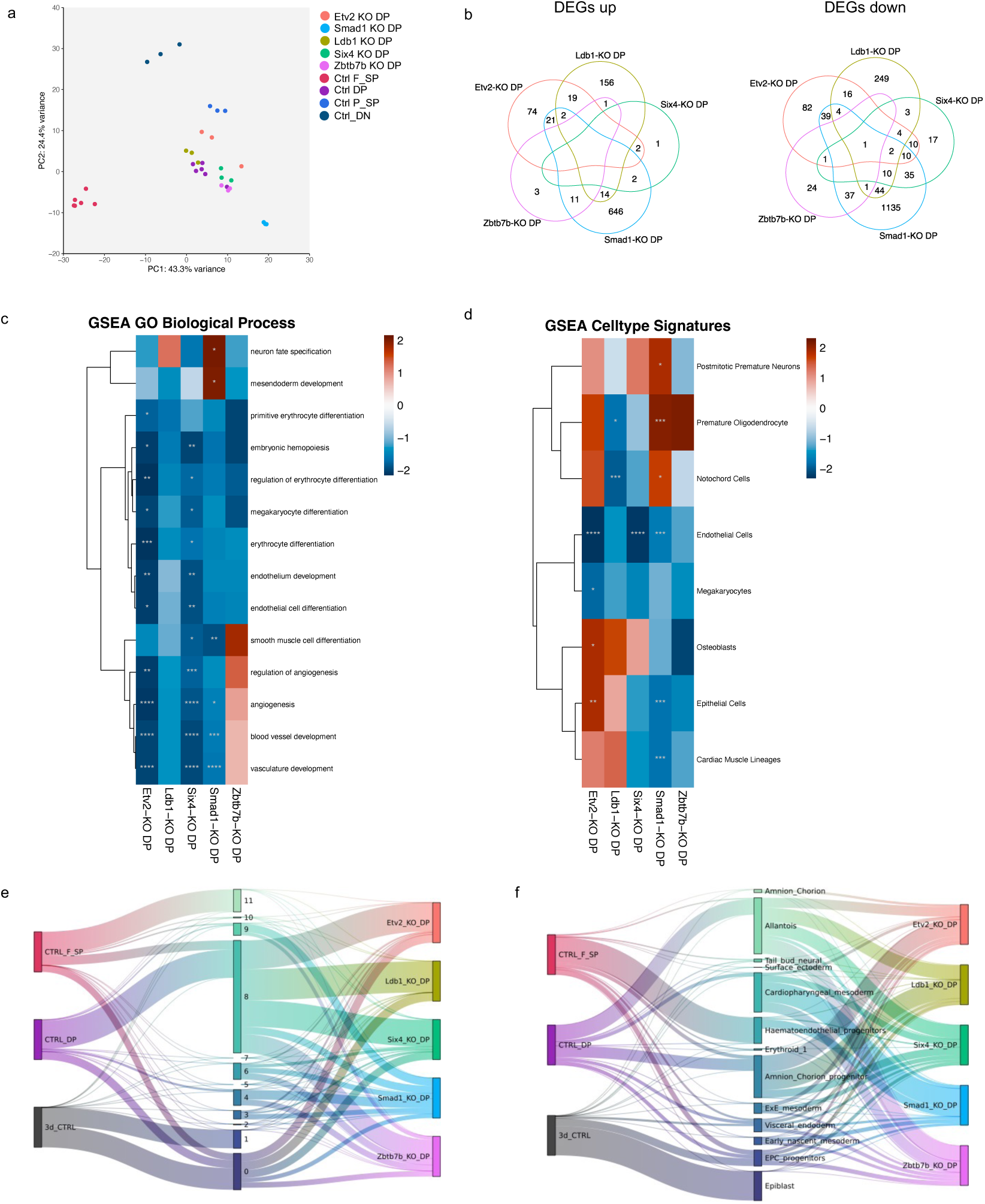
Bulk RNA-seq data analysis of DP cells obtained from EB cultures at day 5 of indicated KOs. **a** Principal component analysis indicates the degree of similarity of DP cells. Each point represents an individually derived KO clone. **b** Venn diagrams depicting the overlaps of (left) upregulated and (right) downregulated differentially expressed genes of indicated KOs in comparison to Ctrls. GSEA of **c** biological processes and **d** cell type signatures of indicated KOs. Sankey plots showing mapping frequencies from deconvolution of bulk RNAseq data with **e** 12 clusters of the mesodermal developmental trajectory calculated from our scRNA-seq dataset of WT cells from EB cultures at day5 and **f** using a publicly available *in vivo* scRNA seq atlas (Mayshar et al.^18^).

**Figure 5:**
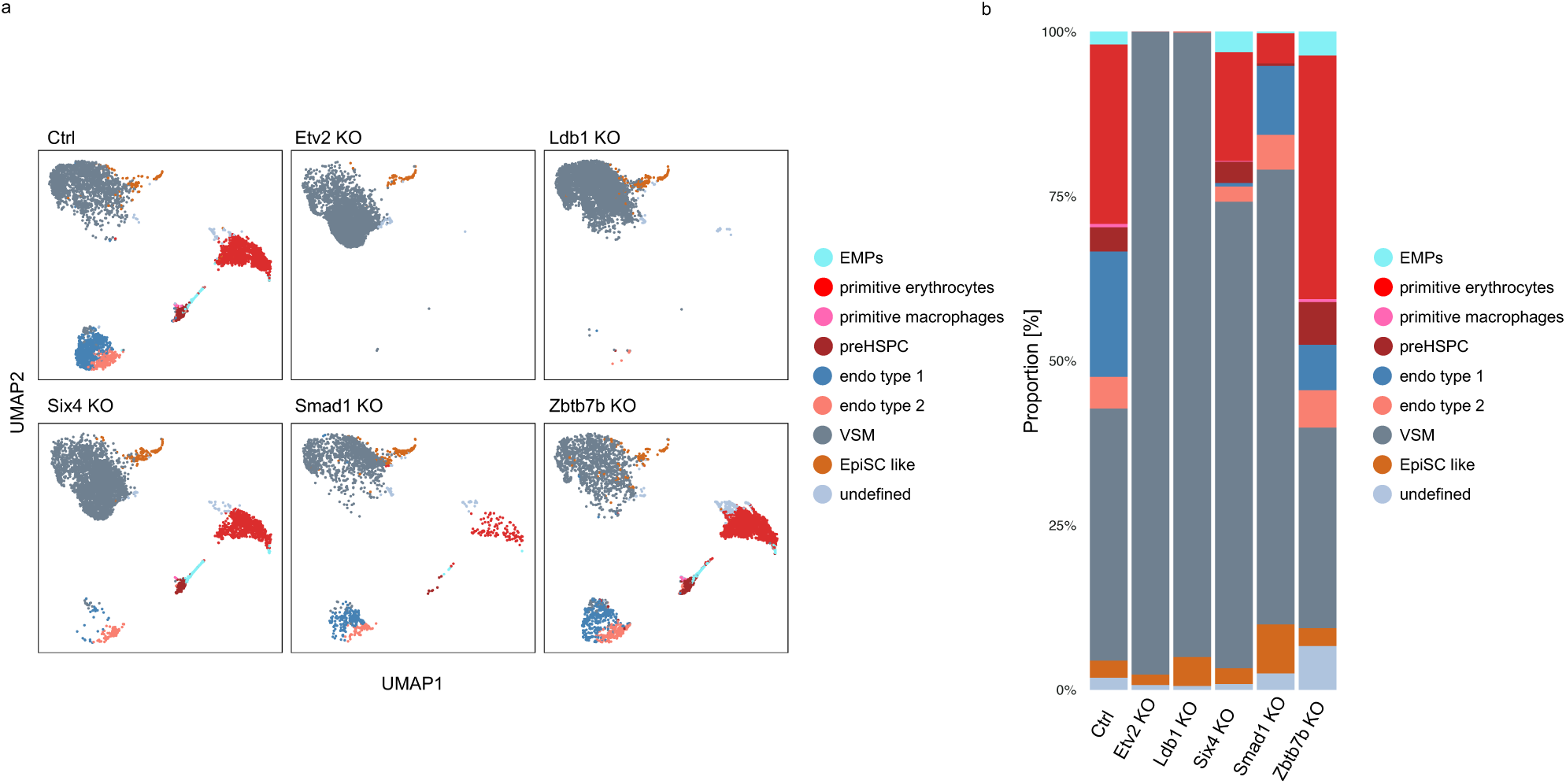
Haematopoietic and endothelial differentiation potential in validated candidate genes. (right) UMAP with overlaid FlowSOM clustering of 50’000 and (left) barchart depicting frequency of cellular subsets identified by FlowSOM clustering in blast culture started from total FLK1^+^ cells of the indicated genotypes.

The Zbtb7b KO DP cells cluster closely with the main group of DPs including the Ctrl and we detected very few DEGs compared to the Ctrl (15 up / 64 down) with minimal overlap to the other KOs (Fig. 4a, Supp. Fig 4a). Additionally, our GSEA analysis detected no significant terms for Zbtb7b KO DPs. We therefore reasoned that the gene expression changes that could lead to an enhanced commitment of Zbtb7b KO cells towards F_SP cells could occur before day 5. We therefore sequenced the EB cultures at day 3 (right at the beginning of mesodermal commitment). We also included the Ctrl and the remaining validated hits (Etv2, Ldb1, Smad1, Six4). We detected few misregulated genes (7 up/ 16 down) in the Zbtb7b KO cells but interestingly, we detected a significant upregulation of the transcription factors Etv2 and Tal1, that are crucial for early blood commitment and could indicate that losing Zbtb7b has an early impact on mesodermal differentiation (Supp.4c). Etv2 and Tal1 were not up-regulated at day3 in any other analysed KO, that showed few DEGs compared to the Ctrls at that time point highlighting that their gene function becomes relevant at the later time point of the differentiation trajectory (Supp Fig.4c).Collectively, our RNAseq analysis clearly underlines the differences of our candidate genes in their impact on gene programs, despite very similar phenotypes (for the depleted candidates) in the EB culture at day 5.

To understand the hierarchy of the candidate TF in F_SP development, we performed deconvolution of the bulk RNA-seq datasets with MuSiC^17^. RNA deconvolution is a powerful tool to infer different cell types of a bulk RNA-seq dataset by comparing it to a reference scRNA-seq atlas. We initially performed a RNA-velocity analysis to assign a latent time to each cell in our scRNAseq dataset of EB differentiation. We assigned 12 clusters along the trajectory, with cluster 0 being at the beginning of the mesodermal developmental trajectory and cluster 11 being the most differentiated one (Supp. Fig. 4d). To confirm the usefulness of the deconvolution analysis we mapped the Ctrl samples to the 12 assigned clusters. As expected, Ctrls isolated at day3 of the differentiation protocol mapped to early clusters (cluster 0 and 1), Ctrl DP cells mapped mainly to the intermediate cluster 8 and Ctrl F_SP mainly to the late cluster 11 (Fig. 4e). Among our KO samples Smad1 KO DP cells showed the strongest difference to the Ctrl DPs and mapped to a wide number of clusters (cluster 0-9) (Fig. 4e). These included earlier clusters (cluster 0-7) but surprisingly also the more advanced cluster 9, which is marked by increased ETV2 expression. While Ldb1-KO exclusively mapped to cluster 8, which corresponds to primitive mesoderm, Etv2- and Six4-KO appear very similar with also small levels of earlier clusters. Interestingly, the Zbtb7b-KO appears reminiscent of Smad1 KO, with a slight expansion of earlier and later clusters (Fig. 4e). In a second step we used a published *in vivo* sc RNA-seq dataset by Mayshar et al.^18^ as a reference. To avoid promiscuous mapping we integrated the *in vivo* dataset with our EB scRNA-seq dataset using Harmony^19^ and only neighbouring cell-types were used for deconvolution in conjunction with a 5% mapping threshold. As a result, the samples were correctly mapped to clusters with a matching developmental stage: Ctrl samples isolated at day 3 mapped mainly to epiblast cells, Ctrl_DP cells to allantois, cardio-pharyngeal mesoderm, amnion-chorion progenitors and ectoplacental cone (EPC) progenitors and Ctrl F_SP to haemato-endothelial progenitors (Fig. 4f). In agreement to the previous analysis Smad1 KO DP cells were the most distinct compared to the Ctrl and other KOs and display a high degree of mapping to cardio-pharyngeal progenitors but also to haemato-endothelial progenitors (Fig. 4f). This recapitulates the deconvolution with our scRNA-seq dataset, as both earlier (cardio-pharyngeal progenitors) and later (haemato-endothelial progenitor) populations, are mapped. Contrary to Etv2 KO and Six4 KO cells, Ldb1 KO cells did not map to extraembryonic mesoderm, which is only found in the former two (Fig. 4f). Surprisingly, but in accordance with the mapping to our EB scRNA-seq reference, Zbtb7b-KO DP cells is reminiscent of Smad1 KO although with lower mapping to haemato-endothelial progenitors (Fig. 4f).

Collectively, the analysis of the RNA-seq data underlines that the different KOs have a diverse effect on the gene expression programs in DP cells. The deconvolution analysis suggests a high degree of heterogeneity within the DN primitive mesoderm, which is a population with a broader developmental potential.

### Haematopoietic and endothelial lineage commitment is differentially regulated by core TF network

To test if our KO lines have indeed a distinct developmental potential as suggested by our RNAseq experiments we proceeded to drive their differentiation towards haematopoietic and endothelial lineages in blast culture. We sorted total FLK1+ cells from EB cultures to include both DP cells and F_SP cells of Ctrl and TF KO cells and analysed their fate conversion with the establish high-dimensional flow cytometry panel. As expected, and in line with the literature, Etv2 KO cells have a complete block towards haematopoietic and endothelial lineage commitment (Ref) (Fig.5a, b). Only VSM and uncommitted EpiSC-like cells form in Etv2 KO cultures. Ldb1 KO cells displayed an equal phenotype to ETV2 KO cells with a complete abrogation of haematopoietic and endothelial cells. Interestingly, and to our surprise, Six4 KO cells were still able to differentiate to haematopoetic lineages including both primitive and definitive subsets (with only a minor decrease) but showed a block towards endothelial lineage, especially endothelial cell type 1 cluster (cKit-) (Fig.5a, b). Smad1 KO cells had a reduce commitment towards both haematopoietic and endothelial lineages. Equally to Six4 KOs, Smad1 KO cells showed mainly a reduction of the endothelial cell type 1 cluster (Fig.5a, b). On the haematopoietic side both primitive and definitive haematopoietic subsets were affected in Smad1 KO cells. Zbtb7b KO cells differentiated towards all lineages with an increase in all haematopoietic lineages and endothelial cell type 2 cluster (Fig.5a, b). This result could be mainly driven by the higher frequency of F_SP cells in the starting culture. Collectively, our data revealed that our experimental approach identified successfully both known and new core TFs with distinct roles in regulating haematopoietic and endothelial linages differentiation, with each TF loss of function leading to the formation of a mesodermal subsets with a defined lineage differentiation bias.

## Discussion

To control cellular differentiation, TFs and chromatin regulators are needed to establish defined gene expression programmes. Early blood differentiation is poorly explored, and we have a limited understanding which molecular regulators impact on the initial steps of blood commitment. In our study, we applied large-scale targeted CRISPR-Cas9 KO screens to investigate the factors governing embryonic blood formation. Our analysis focused on the transition from primitive mesoderm, characterized by a broad developmental potential, to haemato-endothelial mesoderm, the precursor of blood and endothelial lineage. We combine this with an ESC-based differentiation model that faithfully recapitulates yolk sac haematopoiesis^8,9,20^. By using well-defined surface markers including FLK1, PDGFRα, CD41 and VE-cadherin we can pinpoint the developmental subsets of mesodermal and haemato-endothelial origins, that are being formed during the differentiation of pluripotent stem cells towards haematopoietic progenitors. Our scRNA-seq analysis of the *in vitro* generated cells revealed their similarity to cells being formed *in vivo* in the murine yolk sac at E7.5–E8.5. These results underlie the validity of our ESC differentiation system to model yolk sac haematopoiesis and enabled us to uncover the role of several previously unknown factors implicated in early blood commitment. Additionally, murine ESC derived haematopoietic differentiation models have been successfully used in the past to elucidate developmental stages and molecular regulations of blood development that were subsequently validated in *in vivo* models^14,20–22^. By modelling early haematopoiesis *ex vivo,* we were able to obtain sufficient cell numbers to allow large-scale genetic screenings with the optimal sgRNA representation of a developmental delicate time window that would have been challenging if not impossible *in vivo*. We confirmed with high statistical ranking the crucial role of the known master TF ETV2 that enables the transition from primitive to haemato-endothelial mesoderm serving as a positive control for the screening set-up. We additionally identified novel TFs which function as drivers (SMAD1, LDB1 and SIX4) or repressor (ZBTB7b) of haemato-endothelial lineage commitment. Interestingly, we did not validate any chromatin regulator impacting this transition. We speculate that some chromatin regulators play a role earlier in the differentiation trajectory, as we detected them among the top hits of depleted genes in transitions prior to primitive to haemato-endothelial mesoderm (undifferentiated è paraxial mesoderm and paraxial mesoderm è primitive mesoderm) some with known functions in lineage differentiation including: *Kdm6b*, *Kdm6a*, *Kmt2d*, *Arid2* and *Brd8.* By *c*hanging to an inducible screening set-up, when the KOs are induced not only at the pluripotent stage (day 0 of the differentiation protocol) but right before mesodermal commitment occurs (e.g. at day 3 of the differentiation protocol) it would be possible to test if they play a role in the transition of primitive to haemato-endothelial mesoderm as well.

Focusing on the identified drivers their phenotypes in EB cultures were surprisingly very similar. The different KO lines displayed a comparable differentiation block towards haemato-endothelial mesoderm whereas no effect on the differentiation of primitive mesoderm was detected. Besides ETV2, with its known function in haemato-endothelial mesoderm development, Ldb1 was previously suggested as a novel player in early haematopoiesis, precisely at the transition of haemato-endothelial mesoderm^23^. Additionally, Smad1 KO ESC have been reported to be arrested at the primitive mesodermal stage^24^. The distinct function of the identified drivers on haemato-endothelial development became apparent upon further differentiation, whereas ETV2 and LDB1 KO cells had a complete block to form blood or endothelia, SMAD1 KO cells formed both albeit at a marked reduced level compared to WT cells. Our data suggest a high degree of heterogeneity within primitive mesodermal cells, which can be scored by driving their further differentiation. By relying on the two surface markers FLK1 and PDGFRα it is possible to enrich for mesodermal lineages and this classification has been used in numerous publications with a risk of oversimplification. Throughout embryonic development the expression of those markers is mutually exclusive in mesoderm populations with Flk1 single positive mesoderm giving rise to extraembryonic and lateral plate mesoderm subsequently forming blood and vasculature and PDGFRα single positive mesoderm or paraxial mesoderm forming muscle and bones^25–27^. Several studies suggest some exceptions from this classical discrimination, where e.g. also PDGFRα single positive cells give rise to the haemato-endothelial lineage under appropriate conditions^25,28,29^. Still, these studies suggest that most mesodermal populations transition through the FLK1 and PDGFRα double positive primitive mesoderm stage before they further differentiate to more committed progenitors by down-regulating or maintaining one or both markers. It is a challenge in the future to define new surface proteins that give a finer resolution on the primitive mesodermal stage and could serve as predictor for their lineage bias. Our transcriptome analysis of WT and KO primitive mesoderm populations further supports the notion that it is a highly heterogenous stage and our identified drivers impact distinctly on the gene expression programme of this stage. ETV2 has a very narrow expression window in the developing embryo and equally in EB differentiation model. Therefore, most DEGs that were detected in ETV2 KO primitive mesoderm were related to the downstream blood development. The identified driver genes, *Ldb1*, *Smad1*, and *Six4* have a wider expression pattern and their KOs lead to a broader effect on the gene expression in primitive mesoderm. SMAD1 is the signal transducer of bone morphogenetic protein (BMP) subgroup of TGFβ molecules. BMP signals are known to be important regulators of early embryonic development and more precisely BMP4 is a crucial cytokine needed for mesodermal development^30^. It is therefore not surprising that primitive mesoderm that forms in Smad1 KO cells had the highest amount of DEGs in our analysed candidates and SMAD1 KO cells form a distinct group in the multi-dimensional scaling analysis. The deconvolution method suggests an early arrest of Smad1 KO cells along the mesodermal trajectory, but surprisingly, some cells are still able to differentiate to primitive mesoderm that gives rise to both haematopoietic and endothelial lineage. Additionally, in human embryonic stem cells a critical role of BMP4-Smad1 axis together with brachyury has been reported which drives mesodermal differentiation while repressing endoderm commitment^31^. Three of our identified factors lead to early embryonic lethality in KO mice, which is expected for key factors of early haematopoiesis: ETV2 KO mice die at ∼E9.5-10.5 (ref^2^), Ldb1 KO mice ∼ E9.5-10 (ref^32^) and Smad1 KO mice ∼E9.5 (ref^33^). Six4 stands out among the identified drives, as it has never been linked to endothelia or haematopoietic development and is not essential for mouse embryogenesis^34^. The Six gene family consist of six members, where Six1 and Six4 expression are almost identical. As both genes are only separated by 100kb it is possible that they are regulated equally^35^. Six1 KO mice, contrarily to Six4 KOs, are embryonic lethal with reported defects in myogenesis and many organs but not the vasculature or haematopoietic system (Xu, 2003). Six1 and Six4 double KOs have an augmented phenotype compared to Six1 single KOs and have severe craniofacial and rib defects, and general muscle hypoplasia^36^. Intriguingly, Six4 KO primitive mesoderm cells are able to form blood lineages but have a marked reduction in endothelial commitment. Endothelial cells that are ckit negative are almost completely missing in Six4 KO cultures. On the haematopoietic side both primitive and definitive subsets are present in Six4 KO cultures at a comparable level to WT. As endothelial cells are critically needed to form blood it is tempting to speculate that Six4 is involved in the differentiation of endothelia to hemogenic endothelia cells (HECs). Further analysis are needed to understand if Six4 expressing endothelial cells are depleted of hemogenic endothelia and if Six4 has the potential to serve as a reporter gene to enrich for bona fide HECs (Six4 neg). Besides genes that are crucial to form haemato-endothelial mesoderm we also identified genes that potentially repress this transition, which are reflected by an enriched sgRNA abundance in the F_SP population in the CRISPR screenings. For enriched sgRNAs we detected fewer genes with a high statistical ranking compared to the depleted/ driver genes, which is also reflected by the lower number of common overlaps among the top 200 hits of the three independent screens. We therefore used only three potential genes for validation but still identified the zinc finger protein Zbtb7b as a repressor of primitive mesoderm differentiation. Zbtb7b, also known as ThPok, functions in T cell development as a master regulator of CD4/CD8 lineage determination in thymus^37,38^ but was not implicated so far in early blood development. Interestingly, it was shown recently that Zbtb7b together with BMP4 regulated the primed-to-naïve transition (PNT) of pluripotent stem cells in a model of EpiSC to ESC conversion^39^. Zbtb7b together with Zbtb7a were reported to facilitate opening of naive pluripotent chromatin loci and the subsequent activation of nearby genes^39^. Puzzlingly, we detected few misregulted genes at day5 of Zbtb7b KO EB cultures compared to WT. At day3 we detected the upregulation of two crucial genes for haematopoietic commitment, *Etv2* and *Tal1*, respectively. This result could indicate that Zbtb7b KO cells are more prone to commit towards F_SP cells compared to WT cultures and that the differentiation towards a haemato-endothelial fate is established earlier. Future research will address if Zbtb7b, similar to the PNT study, is indeed involved in chromatin remodelling during mesodermal differentiation, where Zbtb7b loss of function leads to a chromatin landscape favourable to haemato-endothelial differentiation. In conclusion, our study identified novel factors involved in haemato-endothelial mesodermal differentiation and in shaping the developmental potential of primitive mesodermal cells and suggests a more complex interplay between TFs than previously anticipated.

## Methods

### Cell culture and cell line generation

Mouse embryonic stem cells (HA36CB1, 129×C57BL/6) were cultivated on 0.2% gelatine coated dishes. ESC medium consisted of DMEM supplemented with 15% fetal calf serum, 1x nonessential amino acids, 1 mM L-glutamine, LIF, and 0.001% b-mercaptoethanol.

The KO cell lines were generated by co-transfecting pX330-U6-Chimeric_BB-CBh-hSpCas9 (Addgene 42230) with two sgRNAs (sequences see table below). Ctrl samples were transfected with non-targeting sgRNAs (sgRNA1_GCGTATCTACCCTACCGCCG, sgRNA2_GCGGTTACCGCGAAAACCAT). pRR-Puro recombination reporter^40^(Addgene 65853) was co-transfected and 36 hours after transfection cells were treated with 2 μg/ml puromycin for 36 h. Positive KO clones were validated by Sanger sequencing. Transfections were conducted using Lipofectamine 3000 reagent (Thermo Fisher Scientific) at a 2:1 Lipofectamine/DNA ratio in OptiMEM (Thermo Fisher Scientific).

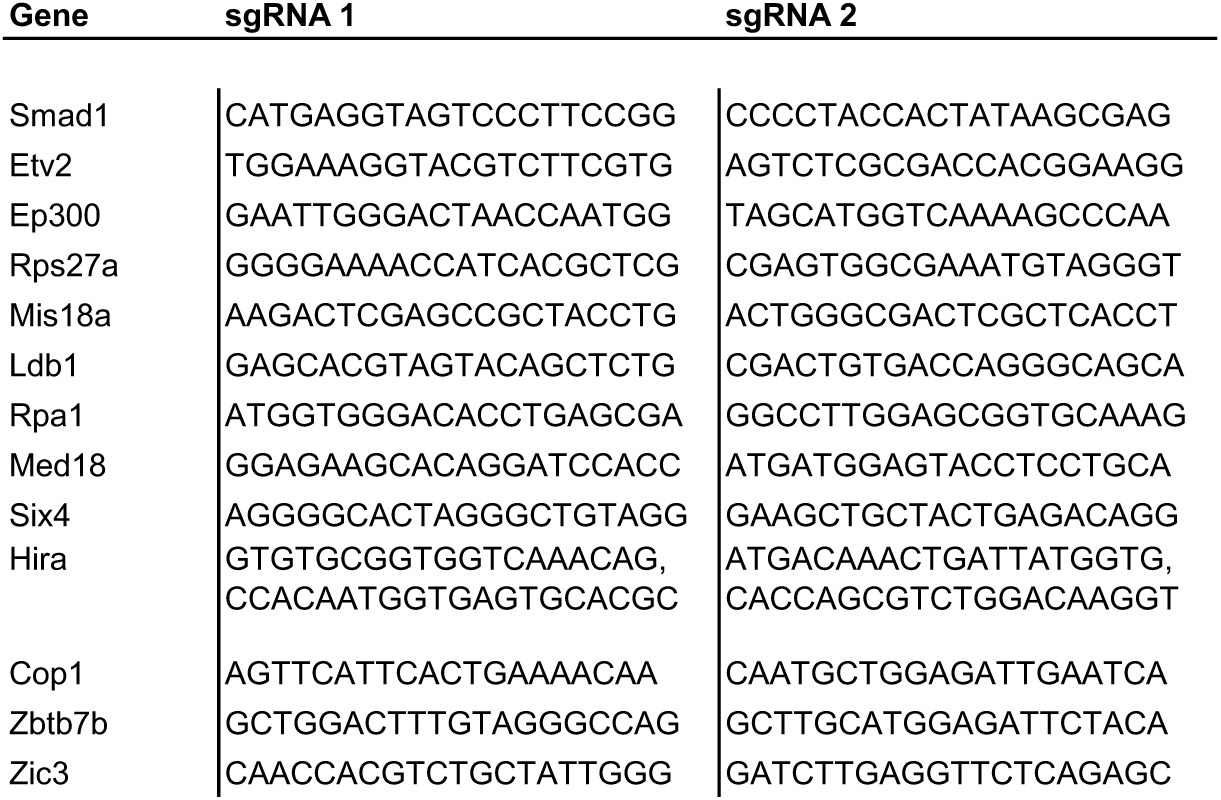

### Embryoid body cultures for mesoderm differentiation

24h prior to the onset of EB cultures, ESCs were transferred on gelatinized plates, using Iscove’s Modified Eagle Medium (IMDM) (Gibco) instead of DMEM. For the generation of embryoid bodie (EB) cultures, ES cells were trypsinized and plated at 10,000 cells/ml in non-adherent 10 cm^2^ petri dishes in EB media containing IMDM supplemented with 1% L-glutamine (Gibco), 10% FBS (Gibco), 0.6% transferrin (Roche, 10652), 50 ug/ml ascorbic acid (Sigma, A4544) and 0.03% monothioglycerol (MTG) (Sigma, M6145). Bmp4 was added from day 0 EB culture and bFGF, activin A and VEGF from day 2.5 (all Peprotech) all at 5 ng/ml. After 5 days in culture EBs were harvested and TrypLETM Express Enzyme (1×) (Gibco, 12605036) was used to generate a single-cell suspension.

### Haemangioblast culture for haematopoietic and endothelial commitment

Prior to blast culture mesodermal populations were FACS sorted (either DP, F_SP or total FLK1+) and 0.5×10^6^ cells were cultured on 0.2% gelatin for 60h in IMDM medium supplemented with 1% Penicillin–Streptomycin, 1% L-glutamine, 15% FBS (Gibco), transferrin (Roche), 0.03% monothioglycerol (MTG) (Sigma, M6145) and 50 mg/μL of ascorbic acid (Sigma, A4544, 5 ng/ml VEGF (Preprotech) and 10 ng/mL IL6 (Preprotech). The cells could then be collected with TrypLE express and cell populations analysed by flow cytometry.

### Flow cytometry

#### EB cultures

Single-cell suspensions were obtained at day 5 of differentiation. For cell-surface staining, cells were incubated for 30 min at 4 °C with a saturating concentration of anti-CD309 (FLK1) monoclonal antibody 1:200 (eBioscience, clone Avas12a1) and anti-CD140a (PDGFRα) monoclonal antibody 1:200 (eBioscience, clone APA5). LIVE/DEAD Fixable Near-IR Dead Cell Stain (L34975, Invitrogen) was used to discriminate cell viability. Samples were acquired using a FACSFortessa (BD Biosciences), and data were analysed using FlowJo software (version 10.7, Tree Star) and visualized with Prism (version 5.0a).

#### Haemangioblast cultures

Single-cell suspensions were obtained at day 2.5 of differentiation Cells were first labeled with Zombie NIR fixable viability dye (BioLegend, 1:500) for the exclusion of dead cells. Surface antigen staining was performed in PBS for 20 minutes at 4°C and washed twice with PBS before acquisition on a Cytek Aurora 5L spectral analyzer. Anti-mouse antibodies used in this study were purchased from BD Biosciences anti-CD117 (BUV395, Clone 2B8), anti-CD43 (BUV496, Clone 1B11), anti-CD45 (BUV563, Clone 30-F11), anti-CD9 (BUV661, Clone KMC8), anti-CD44 (BUV737, Clone IM7), anti-CD31 (BUV805, Clone 390), anti-CD326 (BV480, Clone G8.8), anti-CD93 (BV605, Clone AA4.1), anti-CD41 (BV650, Clone MWReg30), anti-CD14 (FITC, Clone rmC5-3), anti-CD71 (RB780, Clone C2), anti-CD47 (PE-CF594, Clone miap301) or BioLegend including anti-CD64 (BV421, Clone X54-5/7.1), anti-CD16/32 (BV711, Clone 93), anti-CX3CR1 (BV785, Clone SA011F11), anti-CD202b (PE, Clone TEK4), anti-CD34 (PE-Cy5, Clone MEC14.7), anti-CD309 (PE-Cy7, Clone Avas12), anti-Ly6C (Alexa Fluor 700, Clone HK1.4), Anti-Sca-1 (APC-Fire 750, Clone D7), anti-CD11b (APC-Fire 810, Clone M1/70) or ThermoFisher including anti-CD11c (PE-Cy5.5, Clone N418), anti-CD144 (eFluor660, Clone BV13). Spectral unmixing was performed using Spectroflo software (Cytek). Data pre-processing was carried out in FlowJo (BD Biosciences) for singlets and dead cell exclusion. Pre-gated cells were then imported into R studio using R version 4.2.2 using the CATALYST package. The flow cytometry data were then transformed using a hyperbolic arcsine (arcsinh) transformation and percentile normalized to obtain expression values between 0 and 1. This was followed by unifold manifold approximation and projection (UMAP) using the umap implementation of the CATALYST package. Automated clustering and metaclustering of the percentile normalized data were performed with the FlowSOM package. This was followed by expert-guided merging of some clusters based on their median marker expression profile.

### Single cell-sequencing Sort-seq

Viable single cells were FACS sorted into 384-well plates, called cell capture plates, that were ordered from Single Cell Discoveries, a single-cell sequencing service provider based in the Netherlands. Each well of a cell capture plate contains a small 50 nl droplet of barcoded primers and 10 µl of mineral oil (Sigma M8410). After sorting, plates were immediately spun and placed on dry ice. Plates were stored at -80° C. Plates were shipped on dry ice to Single Cell Discoveries, where single-cell RNA sequencing was performed according to an adapted version of the SORT-seq protocol (^41^with primers described in^42^). Cells were heat-lysed at 65° C followed by cDNA synthesis. After second-strand cDNA synthesis, all the barcoded material from one plate was pooled into one library and amplified using in vitro transcription (IVT). Following amplification, library preparation was done following the CEL-Seq2 protocol^43^to prepare a cDNA library for sequencing using TruSeq small RNA primers (Illumina). The DNA library was paired-end sequenced on an Illumina Nextseq™ 500, high output, with a 1×75 bp Illumina kit (read 1: 26 cycles, index read: 6 cycles, read 2: 60 cycles).

#### Data analysis

During sequencing, read 1 was assigned 26 base pairs and was used to identify the Illumina library barcode, cell barcode, and UMI. Read 2 was assigned 60 base pairs and used to map to the reference genome Mus musculus (GRCm38) version 99 with STARSolo 2.7.3a4. Briefly, mapping and generation of count tables were automated using the STARSolo 2.7.3a aligner. Unsupervised clustering and differential gene expression analysis was performed with the Seurat5 R toolkit. Count tables were imported into R, where a singleCellExperiment object was constructed and subsequently filtered based on library size, feature count, spikein count, and mitochondrial count using the isOutlier function from the scater v1.26.1 R package. Following quality control, the dataset was analysed using the Seurat v4.3 R package, using the SCTransform workflow^44^. This included normalisation, scaling, and linear dimensionality reduction.Next, we selected haemato-endothelial cells from the publicly available mouse gastrulation atlas^15^ and integrated them with our in vitro scRNA-seq dataset using Seurat. To complement our integration, we reconstructed the force-directed graph of the haemato-endothelial landscape by incorporating supplementary data from the atlas publication. Within the integrated dataset, we assigned in vitro population labels to the in vivo cells from the atlas based on the k=5 nearest neighbours.For RNA-velocity analysis, the Seurat data was exported and an annotationData object was manually constructed in Python. Subsequently, scvelo v0.2.5 (ref^45^) was applied to it for comprehensive RNA-velocity analysis. The latent time of each cell from the dynamical model was used to calculate an average per cluster, thereby arranging the clusters along the developmental trajectory.

### CRISPR screens in embryoid body cultures

Lentiviral plasmid library targeting TFs and chromatin-regulators (EpiTF library) was kindly provided by Michlits et. al^46^ . Three independent CRISPR screens were performed with a guide representation of ∼ 500x. For lentiviral transduction 60×10^6^ ESCs, that constitutively express Cas9 (ref^16^), were spinfected at 37 °C, 500 x g for 1h using polybrene with a multiplicity of infection (MOI) of 0.2. After spinfection, cells were incubated at 37 °C, 5% CO2 for 48 hours and subsequently FACS sorted based on GFP expression to enrich sgRNA+ cells. Per screen 12×10^6^ FACS sorted sgRNA+ cells were used for EB differentiation. Mesodermal populations of interest were FACS-sorted on day 5 regarding FLK1 and PDGFRα expression. A timepoint zero sample (T0) was collected at the start of the cultures.

### CRISPR screen sequencing library preparation and sequencing

Sampled cells were processed using a commercially available kit for DNA isolation (Qiagen cat. 69504), following the manufacturer’s instructions. DNA concentrations were measured using the Qubit dsDNA BR assay and the entire samples were used for PCR amlifications. In PCR1, sgRNAs were amplified and equipped with adapters using Herculase II Fusion DNA Polymerase (Agilent, #600677) for 25 cycles, utilizing the following primers:

Primer 1: ACACTCTTTCCCTACACGACGCTCTTCCGATCTcttgtggaaaggacgaaacacc Primer 2: caagcagaagacggcatacgagataccgttgatgagtag Following, PCR products were pooled and cleaned up by Qiagen MinElute PCR Purification kit following manufacturer’s instruction (Qiagen cat. 28004) and subsequently size selected with 0.7 x AmpureXP beads, according to the manufacturer’s instructions. DNA concentration of cleaned up PCR products was calculated by Qubit dsDNA HS assay. In PCR2, NEBnext dual indices (NEB #E7600S) were added to the amplicons using NEBNext® Ultra™ II Q5® Master Mix (NEB, cat. M0544S) for 7 cycles. Following, PCR products were pooled and cleaned up by Qiagen MinElute PCR Purification kit following manufacturer’s instruction (Qiagen cat. 28004) and subsequently size selected with 0.7 x AmpureXP beads, according to the manufacturer’s instructions and analysed on an Aligent TapeStation 2000 using High Sensitivity D1000 screen tape. Exact library quantification was carried out via qPCR, using the KAPA Library Quantification Kit (Roche Cat. # 7960140001). Finally, an equimolar pool of all samples was established and sequenced on a NovaSeq 6000 sequencing machine (Illumina Inc., California, USA) with 100 bp single read configuration.

#### CRISPR screens data processing

Basecalling and demultiplexing were performed using bcl2fastq2 v2.20.0.422 (Illumina) and sgRNA-sequences were isolated from the reads using cutadapt v3.1 via hard-clipping. Next, sgRNA-sequences were aligned to the library using bowtie2 v2.3.5 and a countable was assembled. Finally, sgRNA abundances were ranked on the gene-level using the MAGeCK v0.5.9 test module^47^. Rankings were then integrated by calculating the mean rank and p-values were combined by Fisher’s method. Duplicate mean ranks were resolved by additional sub-ranking based on the best rankings across all screens.

### Poly-A RNA-sequencing

For each KO and non-targeting control, 3 individual cell lines were differentiated into EBs. 0.5×10^6^ cells were collected on day 3 and day 5 of EB differentiation. On day 5 mesodermal populations were FACS sorted using FLK and PDGFRα surface expression. Subsequently RNA was isolated using Qiagen RNAeasy kit (Qiagen cat. 74134). The quality of the isolated RNA was determined with a Fragment Analyzer (Agilent, Santa Clara, California, USA). The TruSeq Stranded mRNA protocol (Illumina, Inc, California, USA) was used in the succeeding steps. Briefly, total RNA samples (100-1000 ng) were poly A enriched and then reverse-transcribed into double-stranded cDNA. The cDNA samples were fragmented, end-repaired and adenylated before ligation of TruSeq adapters containing unique dual indices (UDI) for multiplexing. Fragments containing TruSeq adapters on both ends were selectively enriched with PCR. The quality and quantity of the enriched libraries were validated using the Fragment Analyzer. The libraries were normalized to 10 nM in Tris-Cl 10 mM, pH8.5 with 0.1% Tween 20. Single-end 100bp were sequenced on Novaseq 6000 (Illumina, Inc, California, USA) and an average of ∼20 million reads per sample were obtained.

#### Data Analysis

Reads were mapped to the GRCm39.107 genomic reference and count tables generated using STAR-2.7.7 (ref^48^). Count tables were filtered and only genes with a minimum of 10 reads and occurrences in at least three conditions were retained (24155genes). PCA and DEG detection were conducted using DESeq2 v1.38.3 R package^49^and batch correction was performed with the removeBatchEffect function from the limma v3.54.2 R package^50^. For DEG determination, the lfcShrink step was run with apeglm^51^ in DESeq2 and genes with an absolute log2foldchange of at least log2(1.5) and an adjusted p-value of <= 0.05 were selected. Subsequently, GSEA (normalised expression score) and ORA (against expressed genes backgroung) were conducted using clusterProfiler v4.6.2 R package^52^, incorporating gene onthology/biological process and molecular signatures database (MSigDB) annotations (adjusted p-value <= 0.05). Deconvolution of the RNA-seq data was performed using MuSiC v1 R package^17^. To ensure accurate mapping, extensive sanity checks were implemented and only cell types that neighbour our EB scRNA-seq data cells in integrated datasets, created using harmony^19^ (n=20 harmony dims, k=1 nearest neighbour), in conjunction with a 5% mapping threshold, were utilised for deconvolution. Sankey plots were created using d3.js (Ref: https://github.com/d3/d3/releases/tag/v7.8.5, Accessed 2024-01-12).

## Acknowledgements

We thank F. Caiado (University Hospital Zurich) for discussions and advice. We thank members of the Schmolka, Becher and Baubec laboratory for their input and criticism. Furthermore, we thank members of the Functional Genomics Centre Zurich for their genomics support. We thank Single Cell Discoveries for their single-cell sequencing services.

## Author contributions

N.S. and M.T. conceived and designed the study. N.S., M.T., T.W, S.B., and P.Z. developed tools and performed experiments. N.S., M.T., T.W. and I.M. analysed data. A.R.G. gave bioinformatic support. N.S. supervised the research. N.S. wrote the manuscript with input from all authors.

## Supplementary Figures

**Supplementary Figure 1:**
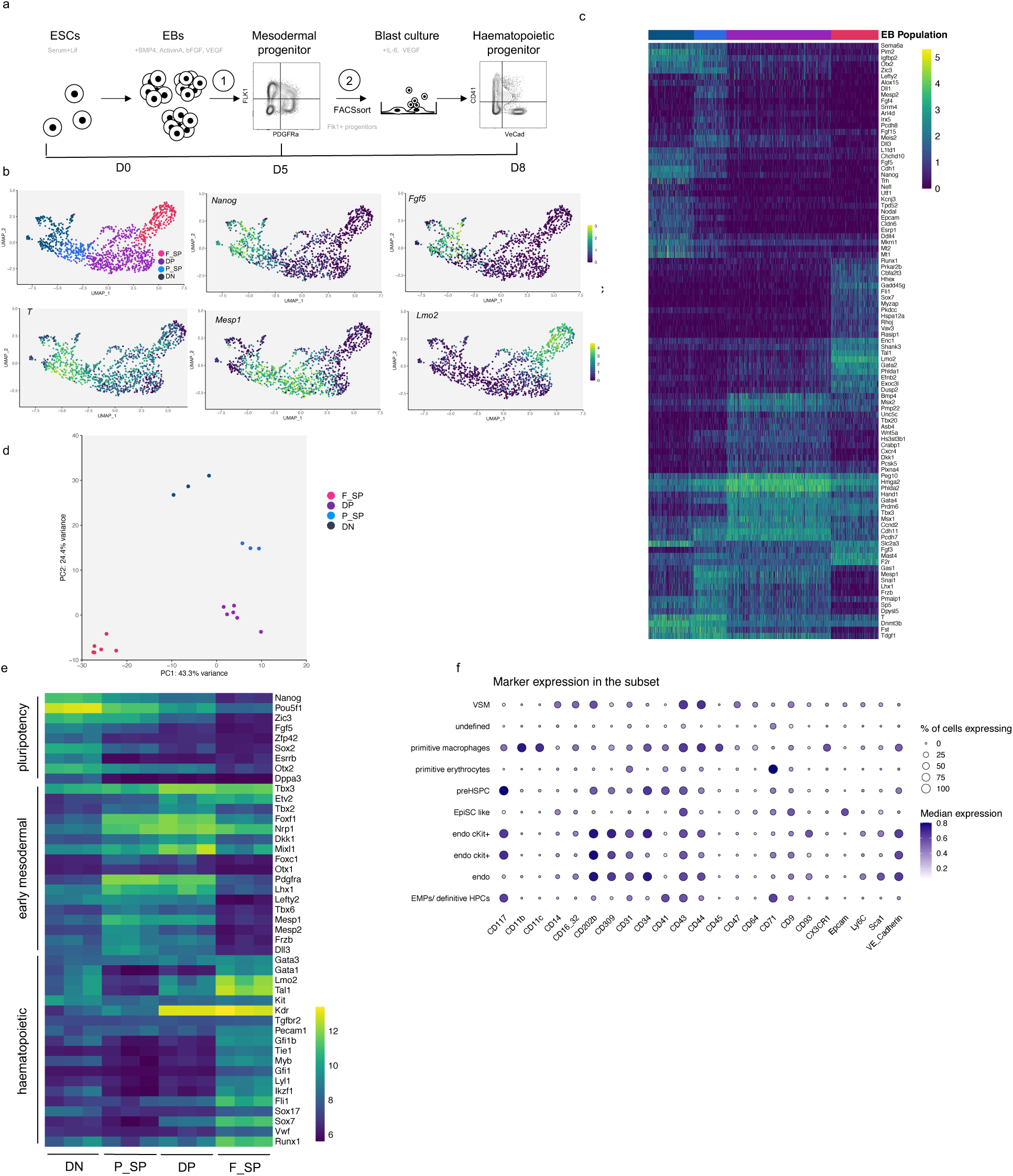
**a** Schematic of ESC-derived haematopoietic differentiation model. **b** Expression of selected mesodermal and haematopoietic signature genes in mesodermal populations analysed by scRNA-seq. **c** Heatmap indicating gene expression by RNA-seq of top 20 differential regulated genes in each mesodermal population. **d** Principal component analysis of bulk RNA-seq data of mesodermal populations obtained from EB cultures at day5. Each point represents an individual culture. **e** Heatmap indicating gene expression by bulk RNA-seq of pluripotency-, mesoderm- and haematopoiesis-associated genes in the analysed mesodermal populations (VST values). **f** Expression matrix of markers used in high-dimensional flow cytometry analysis in DP and F_SP cells.

**Supplementary Figure 2:**
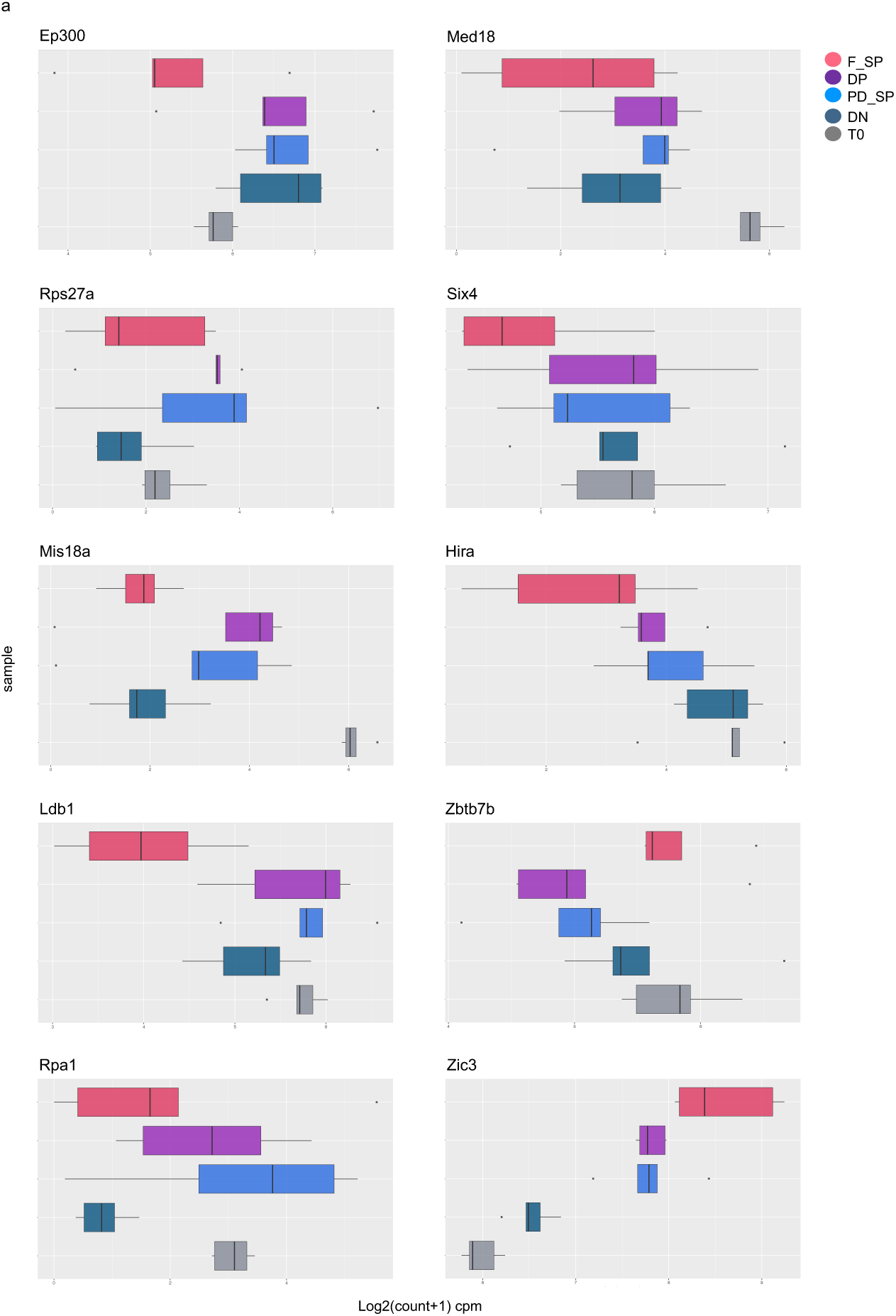
Count per million (CPM) values of sgRNA abundance at T0 and mesodermal populations of: *Smad1*, *Etv2*, *Cop1* and *Zbtb7b*.

**Supplementary Figure 3:**
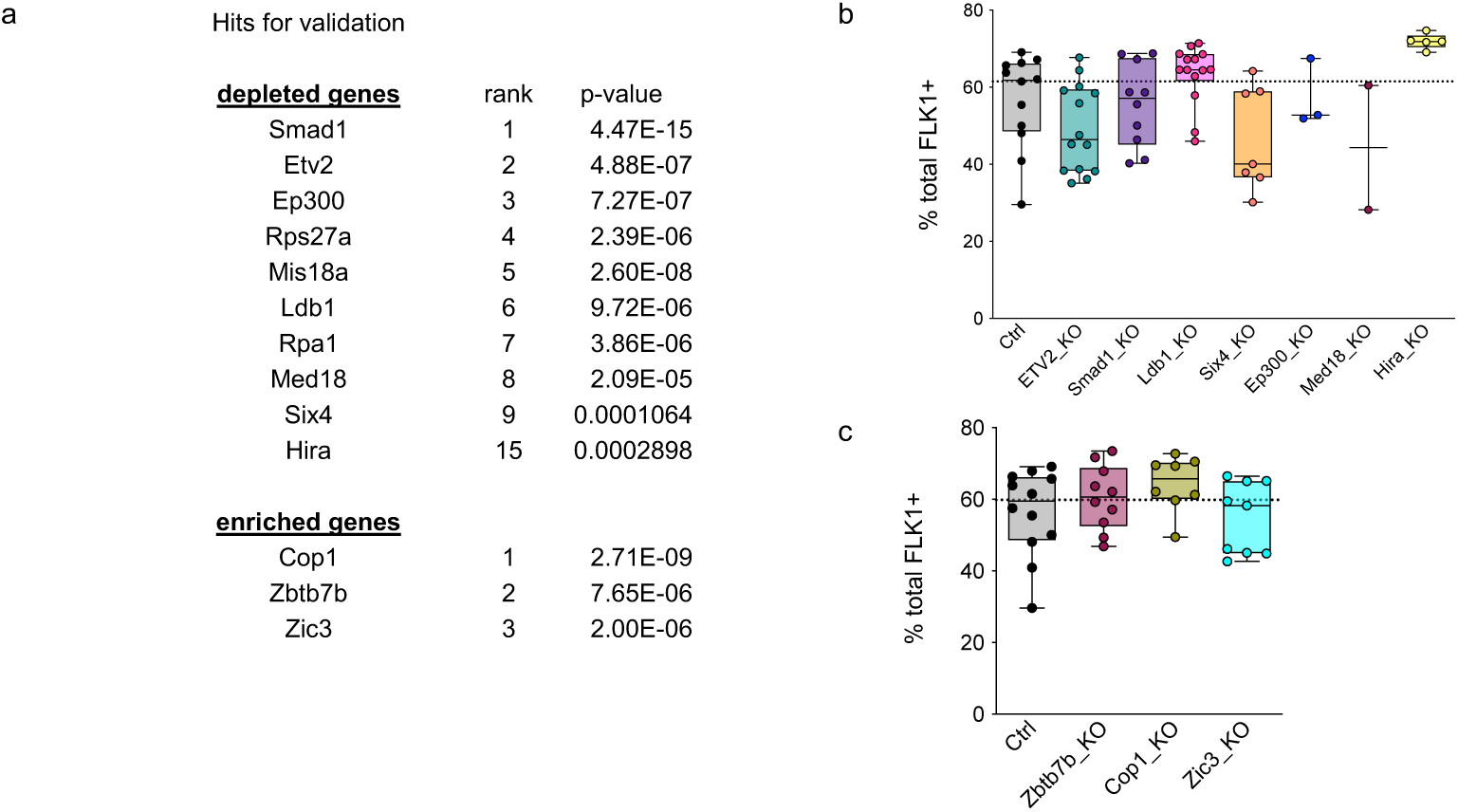
**a** Table of hits that were used for validation experiments of depleted and enriched genes. **b, c** percentage of total FLK cells in EB cultures of indicated KOs. Each data point (in b,c) represents an individually generated cell line. For each cell line 3 independent clones were analysed, n = 4 biological replicates. Error bars represent mean +/− SD. p-values were calculated using a one-way ANOVA multiple comparisons analysis.

**Supplementary Figure 4:**
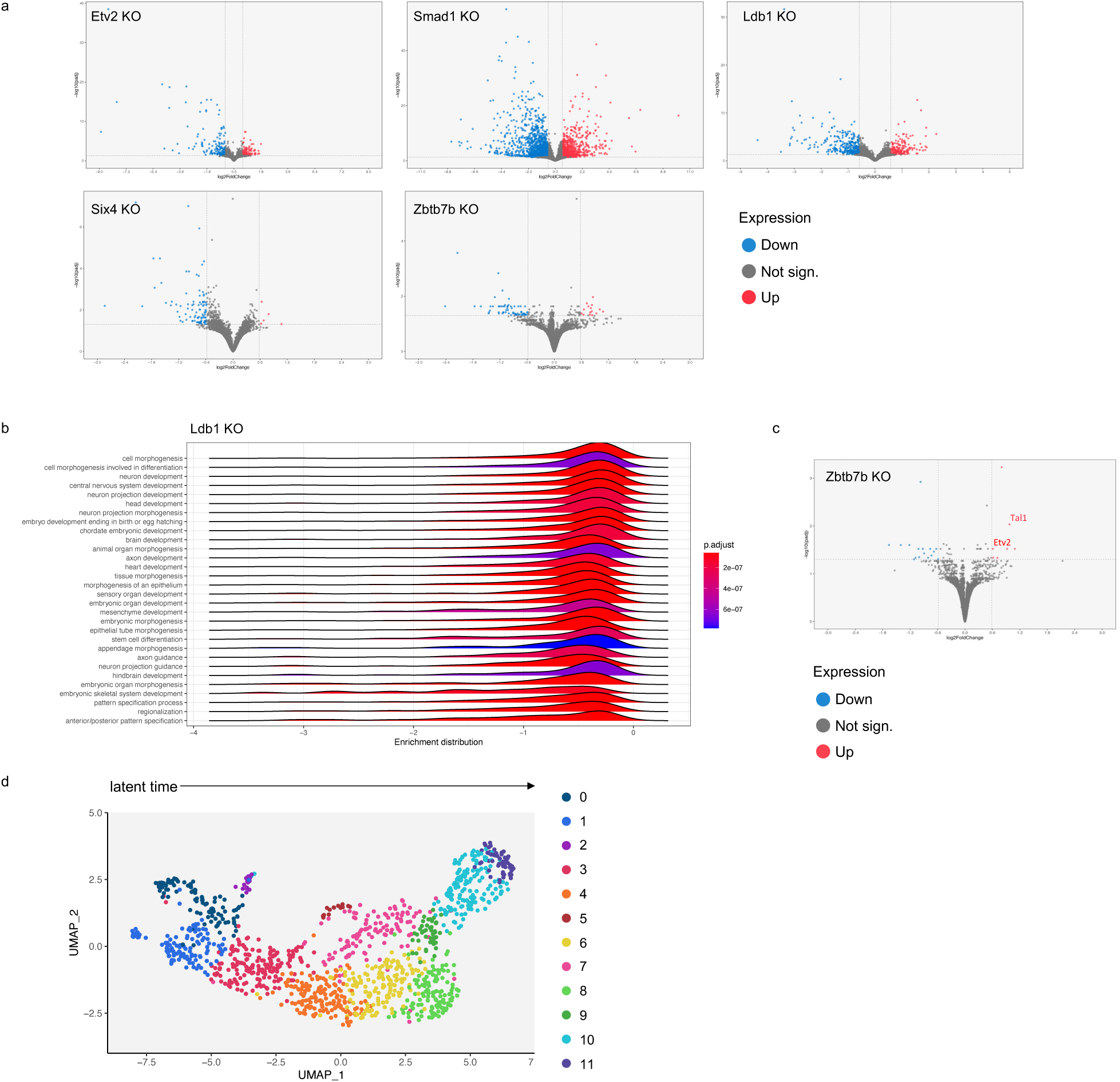
**a** Volcano-plots showing differential gene expression of DP cells of EB cultures at day 5 of indicated KOs vs. Ctrl. Red and blue dots indicate genes with significant changes in gene expression (DeSeq2, |log2FC| >= log2(1.5) and adjusted p-value <= 0.05). **b** GSEA for LBD1 KO DP cells obtained from EB cultures at day 5. **c** Volcano-plot showing differential gene expression of DP cells of EB cultures at day 3 of Zbtb7b vs. Ctrl. Red and blue dots indicate genes with significant changes in gene expression (Deseq2, |log2FC| >= log2(1.5) and adjusted p-value <= 0.05). **d** UMAP projection of RNA velocity analysis assigning 12 clusters along the EB trajectory.

## References

1. Ferdous, A. et al. Nkx2-5 transactivates the Ets-related protein 71 gene and specifies an endothelial/endocardial fate in the developing embryo. Proc Natl Acad Sci U S A 106, 814– 819 (2009).

2. Lee, D. et al. ER71 acts downstream of BMP, Notch, and Wnt signaling in blood and vessel progenitor specification. Cell Stem Cell 2, 497–507 (2008).

3. De Val, S. et al. Combinatorial Regulation of Endothelial Gene Expression by Ets and Fork-head Transcription Factors. Cell 135, 1053–1064 (2008).

4. Ginsberg, M. et al. Efficient direct reprogramming of mature amniotic cells into endothelial cells by ETS factors and TGFβ suppression. Cell 151, 559–575 (2012).

5. Sinha, T. et al. Differential Etv2 threshold requirement for endothelial and erythropoietic development. Cell Rep 39, (2022).

6. Tosic, J. et al. Eomes and Brachyury control pluripotency exit and germ-layer segregation by changing the chromatin state. Nat Cell Biol 21, 1518–1531 (2019).

7. Lancrin, C. et al. The haemangioblast generates haematopoietic cells through a haemogenic endothelium stage. Nature (2009) doi:10.1038/nature07679.

8. Lancrin, C. et al. Blood cell generation from the hemangioblast. Journal of Molecular Medicine vol. 88 167–172 Preprint at 10.1007/s00109-009-0554-0 (2010).

9. Fehling, H. J. et al. Tracking mesoderm induction and its specification to the hemangioblast during embryonic stem cell differentiation. Development 130, 4217–27 (2003).

10. Kataoka, H. et al. Etv2/ER71 induces vascular mesoderm from Flk1 +PDGFRα + primitive mesoderm. Blood 118, 6975–6986 (2011).

11. Kattman, S. J., Huber, T. L. & Keller, G. M. M. Multipotent Flk-1+ Cardiovascular Progenitor Cells Give Rise to the Cardiomyocyte, Endothelial, and Vascular Smooth Muscle Lineages. Dev Cell 11, 723–732 (2006).

12. Choi, K., Kennedy, M., Kazarov, A., Papadimitriou, J. C. & Keller, G. A common precursor for hematopoietic and endothelial cells. Development 125, 725–32 (1998).

13. Ema, M. et al. Combinatorial effects of Flk1 and Tal1 on vascular and hematopoietic development in the mouse. Genes Dev 17, 380–393 (2003).

14. Lancrin, C. et al. GFI1 and GFI1B control the loss of endothelial identity of hemogenic endothelium during hematopoietic commitment. Blood (2012) doi:10.1182/blood-2011-10-386094.

15. Pijuan-Sala, B. et al. A single-cell molecular map of mouse gastrulation and early organogenesis. Nature 566, 490–495 (2019).

16. Butz, S. et al. DNA sequence and chromatin modifiers cooperate to confer epigenetic bistability at imprinting control regions. Nat Genet 54, 1702–1710 (2022).

17. Wang, X., Park, J., Susztak, K., Zhang, N. R. & Li, M. Bulk tissue cell type deconvolution with multi-subject single-cell expression reference. Nature Communications 2019 10:1 10, 1–9 (2019).

18. Mayshar, Y. et al. Time-aligned hourglass gastrulation models in rabbit and mouse. Cell 186, 2610–2627.e18 (2023).

19. Korsunsky, I. et al. Fast, sensitive and accurate integration of single-cell data with Harmony. Nat Methods 16, 1289–1296 (2019).

20. Lancrin, C. et al. The haemangioblast generates haematopoietic cells through a haemogenic endothelium stage. Nature 457, 892–895 (2009).

21. Eilken, H. M., Nishikawa, S. I. & Schroeder, T. Continuous single-cell imaging of blood generation from haemogenic endothelium. Nature 457, 896–900 (2009).

22. Harland, L. T. G. et al. The T-box transcription factor Eomesodermin governs haemogenic competence of yolk sac mesodermal progenitors. Nat Cell Biol 23, 61–74 (2021).

23. Mylona, A. et al. Genome-wide analysis shows that Ldb1 controls essential hematopoietic genes/pathways in mouse early development and reveals novel players in hematopoiesis. Blood 121, 2902–2913 (2013).

24. Cook, B. D., Liu, S. & Evans, T. Smad1 signaling restricts hematopoietic potential after promoting hemangioblast commitment. Blood 117, 6489–6497 (2011).

25. Ding, G., Tanaka, Y., Hayashi, M., Nishikawa, S. I. & Kataoka, H. PDGF Receptor Alpha+ Mesoderm Contributes to Endothelial and Hematopoietic Cells in Mice. Developmental Dynamics 242, 254–268 (2013).

26. Kataoka, H. et al. Expressions of PDGF receptor alpha, c-Kit and Flk1 genes clustering in mouse chromosome 5 define distinct subsets of nascent mesodermal cells. Dev Growth Differ 39, 729–740 (1997).

27. Shalaby, F. et al. A requirement for Flk1 in primitive and definitive hematopoiesis and vasculogenesis. Cell 89, 981–990 (1997).

28. Motoike, T., Markham, D. W., Rossant, J. & Sato, T. N. Evidence for novel fate of Flk1+ progenitor: contribution to muscle lineage. Genesis 35, 153–159 (2003).

29. Chan, S. S. K. et al. Mesp1 patterns mesoderm into cardiac, hematopoietic, or skeletal myogenic progenitors in a context-dependent manner. Cell Stem Cell 12, 587–601 (2013).

30. Winnier, G., Blessing, M., Labosky, P. A. & Hogan, B. L. M. Bone morphogenetic protein-4 is required for mesoderm formation and patterning in the mouse. Genes Dev 9, 2105–2116 (1995).

31. Faial, T. et al. Brachyury and SMAD signalling collaboratively orchestrate distinct mesoderm and endoderm gene regulatory networks in differentiating human embryonic stem cells. Development 142, 2121–2135 (2015).

32. Mukhopadhyay, M. et al. Functional ablation of the mouse Ldb1 gene results in severe patterning defects during gastrulation. Development 130, 495–505 (2003).

33. Tremblay, K. D., Dunn, N. R. & Robertson, E. J. Mouse embryos lacking Smad1 signals display defects in extra-embryonic tissues and germ cell formation. Development 128, 3609–3621 (2001).

34. Ozaki, H. et al. Six4, a putative myogenin gene regulator, is not essential for mouse embryonal development. Mol Cell Biol 21, 3343–3350 (2001).

35. Boucher, C. A., Carey, N., Edwards, Y. H., Siciliano, M. J. & Johnson, K. J. Cloning of the human SIX1 gene and its assignment to chromosome 14. Genomics 33, 140–142 (1996).

36. Grifone, R. et al. Six1 and Six4 homeoproteins are required for Pax3 and Mrf expression during myogenesis in the mouse embryo. Development 132, 2235–2249 (2005).

37. Sun, G. et al. The zinc finger protein cKrox directs CD4 lineage differentiation during intrathymic T cell positive selection. Nat Immunol 6, 373–381 (2005).

38. He, X. et al. The zinc finger transcription factor Th-POK regulates CD4 versus CD8 T-cell lineage commitment. Nature 433, 826–833 (2005).

39. Yu, S. et al. BMP4 resets mouse epiblast stem cells to naive pluripotency through ZBTB7A/B-mediated chromatin remodelling. Nat Cell Biol 22, 651–662 (2020).

40. Flemr, M. & Bühler, M. Single-step generation of conditional knockout mouse embryonic stem cells. Cell Rep 12, 709–716 (2015).

41. Muraro, M. J. et al. A Single-Cell Transcriptome Atlas of the Human Pancreas. Cell Syst 3, 385–394.e3 (2016).

42. Van Den Brink, S. C., et al. Single-cell sequencing reveals dissociation-induced gene expression in tissue subpopulations. Nat Methods 14, 935–936 (2017).

43. Hashimshony, T. et al. CEL-Seq2: Sensitive highly-multiplexed single-cell RNA-Seq. Genome Biol 17, 77 (2016).

44. Hafemeister, C. & Satija, R. Normalization and variance stabilization of single-cell RNA-seq data using regularized negative binomial regression. Genome Biol 20, 1–15 (2019).

45. Bergen, V., Lange, M., Peidli, S., Wolf, F. A. & Theis, F. J. Generalizing RNA velocity to transient cell states through dynamical modeling. Nature Biotechnology 2020 38:12 38, 1408–1414 (2020).

46. Michlits, G. et al. CRISPR-UMI: single-cell lineage tracing of pooled CRISPR–Cas9 screens. Nature Methods 2017 14:12 14, 1191–1197 (2017).

47. Li, W. et al. MAGeCK enables robust identification of essential genes from genome-scale CRISPR/Cas9 knockout screens. Genome Biol 15, 554 (2014).

48. Dobin, A. et al. STAR: ultrafast universal RNA-seq aligner. Bioinformatics 29, 15–21 (2013).

49. Love, M. I., Huber, W. & Anders, S. Moderated estimation of fold change and dispersion for RNA-seq data with DESeq2. Genome Biol 15, 550 (2014).

50. Ritchie, M. E. et al. A comparison of background correction methods for two-colour micro-arrays. Bioinformatics 23, 2700–7 (2007).

51. Zhu, A., Ibrahim, J. G. & Love, M. I. Heavy-tailed prior distributions for sequence count data: removing the noise and preserving large differences. Bioinformatics 35, 2084–2092 (2019).

52. Wu, T. et al. clusterProfiler 4.0: A universal enrichment tool for interpreting omics data. Innovation (Cambridge (Mass.)) 2, (2021).

